# Dynamic Causal Modeling of Neural Responses to an Orofacial Pneumotactile Velocity Array

**DOI:** 10.1101/2021.04.12.439574

**Authors:** Yingying Wang, Rebecca Custead, Hyuntaek Oh, Steven M. Barlow

## Abstract

The effective connectivity of neuronal networks during orofacial pneumotactile stimulation with different velocities is still unknown. The present study aims to characterize the effectivity connectivity elicited by three different saltatory velocities (5, 25, and 65 cm/s) over the lower face using dynamic causal modeling on functional magnetic resonance imaging data of twenty neurotypical adults. Our results revealed the contralateral SI and SII as the most likely sources of the driving inputs within the sensorimotor network for the pneumotactile stimuli, suggesting parallel processing of the orofacial pneumotactile stimuli. The 25 cm/s pneumotactile stimuli modulated forward interhemispheric connection from the contralateral SII to the ipsilateral SII, suggesting a serial interhemispheric connection between the bilateral SII. Moreover, the velocity pneumotactile stimuli influenced the contralateral M1 through both contralateral SI and SII, indicating that passive pneumotactile stimulation may positively impact motor function rehabilitation. Furthermore, the slow velocity 5 cm/s pneumotactile stimuli modulated both forward and backward connections between the right cerebellar lobule VI and the contralateral left SI, SII, and M1, while the medium velocity 25 cm/s pneumotactile stimuli modulated both forward and backward connections between the right cerebellar lobule VI and the contralateral left SI and M1. Our findings suggest that the right cerebellar lobule VI plays a role in the sensorimotor network through feedforward and feedback neuronal pathways.

## 1. INTRODUCTION

Functional brain imaging studies have provided insights into the human sensorimotor network. The primary (SI) and secondary (SII) somatosensory cortices, and the primary motor cortex (M1) are the core brain regions within the sensorimotor network (Ackerley et al., 2012; Custead, Oh, Wang, & Barlow, 2017; Grodd, Hulsmann, Lotze, Wildgruber, & Erb, 2001; Oh, Custead, Wang, & Barlow, 2017). The functional representations of moving tactile stimulation have primarily used electric stimulation or passive touch on the glabrous hand (Ackerley et al., 2012; Lin & Kajola, 2003; Oh et al., 2017). The contralateral SI and bilateral SII have been activated during various types of touch (Ackerley et al., 2012; Disbrow, Roberts, Poeppel, & Krubitzer, 2001; Francis et al., 2000; Ionta, Martuzzi, Salomon, & Blanke, 2014; Ruben et al., 2001). The contralateral M1 has been involved during passive touch or air pressure pule stimulation to the hands’ glabrous skin (Ackerley et al., 2012; Francis et al., 2000; Oh et al., 2017). For the face, there is scanty evidence on how moving tactile stimulation is processed in the brain. Our group previously identified a putative neuronal somatosensory network with bilateral SI, left M1, and the right lobule VI using a univariate generalized linear model (GLM) on functional magnetic resonance imaging (fMRI) data of 20 neurotypical adults when perceiving moving stimulation on the right lower face (Custead et al., 2017). A subsequent functional connectivity (FC) analysis on the same data revealed that the medium velocity (25 cm/s) tactile stimulation evoked stronger FC in the ipsilateral cortical regions than the low velocity (5 cm/s) (Wang, Sibaii, Custead, Oh, & Barlow, 2020). Nevertheless, there is still much to learn about the complex neuronal networks involved in orofacial pneumotactile velocity processing.

The serial processing theory suggests that somatosensory stimulation flows predominantly from the contralateral thalamus to the contralateral SI and then to the contralateral SII. The contralateral SI receives the driving inputs from tactile stimuli and initiates the higher-order processing of the spatiotemporal information about the tactile stimuli (Disbrow et al., 2001; Lundblad, Olausson, Hermansson, & Wasling, 2011; Norrsell & Olausson, 1994). Animal studies using electrophysiological and anatomical tracing approaches found extensive cortico-cortical projections between SI and SII (Burton & Carlson, 1986; Friedman, Jones, & Burton, 1980; Pons & Kaas, 1986). In humans, a small number of studies using dynamic causal modeling (DCM) with functional magnetic resonance imaging (fMRI) data have suggested that both innocuous and noxious tactile stimuli are processed in serial mode (from SI to SII) (Kalberlah, Villringer, & Pleger, 2013; Khoshnejad, Piché, Saleh, Duncan, & Rainville, 2014). In contrast, the parallel processing theory proposes that somatosensory stimulation is directly transmitted to both contralateral SI and SII. Animal studies have reported that direct thalamocortical inputs to SII bypassing SI (Rowe, Turman, Murray, & Zhang, 1996; Turman, Ferrington, Ghosh, Morley, & Rowe, 1992; Zhang et al., 2001; Zhang et al., 1996) and direct projections from thalamic nuclei (i.e., the ventral posterior nucleus, the ventral posterior inferior nucleus, etc.) to both SI and SII (Jones, 1998; Krubitzer & Kaas, 1992). Several human studies supported the parallel processing theory using various imaging approaches (Klingner et al., 2015; Liang, Mouraux, & Iannetti, 2011; Raij et al., 2008; Song et al., 2021). A multimodal imaging study suggested that parallel inputs to both SI and SII facilitate long-distance cortico-cortical connections using magnetoencephalography (MEG) and single-pulse transcranial magnetic stimulation (TMS) with electroencephalography (EEG) during electrical stimuli to the dominant hand’s median nerve (Raij et al., 2008). Liang et al. (2011) identified that the neural activities elicited by both innocuous and noxious tactile stimuli are best explained by DCM models where the fMRI responses in both SI and SII depend on direct inputs from the thalamus. Moreover, DCM studies of both MEG and fMRI data supported the parallel processing theory for both nociceptive and tactile information processing (Klingner et al., 2015; Song et al., 2021). Another DCM study of fMRI data revealed the coexistence of the serial and parallel modes between SI and SII during pressure stimulation (Chung et al., 2014). Taken together, whether tactile stimuli are processed in serial or parallel mode or both modes in humans remains a topic of debate. Therefore, the present study will examine how orofacial pneumotactile stimuli are processed (serial or parallel or both modes) and whether the neuronal mechanism of tactile processing depends on different velocity tactile inputs.

The bilateral activation of SII elicited by unilateral somatosensory stimuli (Custead et al., 2017; Forss, Salmelin, & Hari, 1994; Hari et al., 1993; Hoechstetter et al., 2001) and the existence of dense transcallosal fibers connecting the contralateral and ipsilateral SII (Jones & Powell, 1969b; Pandya & Vignolo, 1968; Picard, Lepore, Ptito, & Guillemot, 1990) suggest that there are interhemispheric connections between the bilateral SII. In addition, the size of the corpus callosum was positively correlated with the peak amplitude of the ipsilateral SII during innocuous electrical stimuli to the right index finger (Stancak et al., 2002). Patients with callosotomy failed to show ipsilateral SII activation during unilateral tactile stimulation (Fabri et al., 1999). Therefore, both structural and functional interhemispheric connections exist between the bilateral SII. The present study will identify how pneumotactile information flows from the contralateral SII to the ipsilateral SII (i.e., via forward, backward, or both connections) in neurotypical adults, providing important evidence on typical interhemispheric connections between bilateral SII. Brain injury caused by stroke or traumatic brain injury can induce interhemispheric changes and change activity in both affected hemisphere via transcallosal inhibition and unaffected hemisphere via transcallosal disinhibition (Bannister, Crewther, Gavrilescu, & Carey, 2015; Cramer & Crafton, 2006; Mohajerani, Aminoltejari, & Murphy, 2011; Murase, Duque, Mazzocchio, & Cohen, 2004; Pellegrino et al., 2012). Functional connectivity studies have shown that resting-state interhemispheric functional connectivity is associated with stroke recovery (Carter et al., 2010; Compston, 2011). This study will determine effective interhemispheric connectivity between the bilateral SII, which can be useful for evaluating functional brain reorganization after brain injury.

The cross-modality plasticity theory suggested that passive somatosensory stimuli could elicit neuronal responses to improve motor function (Ladda et al., 2014; Nasir, Darainy, & Ostry, 2013; Pearson, 2000; Sanes & Donoghue, 2000). The face sensorimotor network is essential for speech production, sucking, and swallowing, and the sensorimotor integration is critical for motor control and learning (Barlow & Estep, 2006; Barlow & Stumm, 2010; Sessle et al., 2007; Sessle et al., 2005; Smith, 2016). Little is known about how orofacial pneumotactile information flows within the SI, SII, and M1 network. Passive motor and sensory stimulation of hands and feet elicited equal activation levels in the sensorimotor cortex as the active motor tasks (Blatow et al., 2011). High-frequency passive repetitive sensory stimulation, utilizing Hebbian learning principles, has been successfully used to treat chronic stroke patients and improved sensorimotor functions without the need for active participation (Ahn, Lee, & Hwang, 2016b; Chen et al., 2018a; Conforto, Cohen, Dos Santos, Scaff, & Marie, 2007; Powell, Pandyan, Granat, Cameron, & Stott, 1999; Smith, Dinse, Kalisch, Johnson, & Walker-Batson, 2009). If passive orofacial pneumotactile stimulation effectively elicits changes in M1 and positively impact motor function, patients unable to perform active movements right after brain injury (e.g., due to stroke, traumatic brain injury, etc.) could benefit from early interventions with passive sensory and motor stimulation. Moreover, a combination of passive sensory stimulation and active movement training might induce more neural plasticity by integrating sensory and motor systems compared to either sensory or motor modality alone. Thus, understanding the neural mechanism underlying how passive orofacial pneumotactile processing in the present study is critical for uncovering the impacts of somatosensory stimulation on motor function rehabilitation. The effective connectivity within the sensory and motor system revealed by DCM will yield the neural pathways supporting sensory-motor integration.

The cerebellum is fundamentally recognized to be involved in motor control and motor learning (Albus, 1971; Marr, 1969), and growing evidence also suggests its role in the processing of cognition and emotion (Buckner, 2013; D’angelo & Casali, 2012; Schmahmann & Caplan, 2006). Through the cerebellar peduncles, all cerebellar nuclei are interconnected with the rest of the brain. The dentate nucleus, connected to thalamic nuclei and sensorimotor regions through the superior peduncle (Dum & Strick, 2003; Tellmann et al., 2015), has been involved in speech or cognitive learning (Thurling et al., 2011). Recent evidence has supported that the cerebellum can be subdivided into several specialized functional regions (Witter & De Zeeuw, 2015). Previous work has demonstrated that the right lobule VI was a part of the sensorimotor somatotopic representations for the face (Custead et al., 2017; Grodd et al., 2001; Wang et al., 2020). The effective connectivity of the cortico-cerebral networks elicited by passive orofacial pneumotactile stimulation has not yet been studied, which will provide unique insights into the feedforward and feedback pathways of tactile processing in the cortico-cerebral networks.

Our previous study identified similarities and differences of functional connectivity in the sensorimotor system during orofacial pneumotactile stimuli of different velocities (5, 25, 65 cm/s) and shed light on the functional networks encoding the orofacial pneumotactile perception of velocity (Wang et al., 2020). However, the functional connectivity is limited to undirected connections among regions, and the causal relationships within the sensorimotor network cannot be examined. The directed causal influences among neural populations are defined as effective connectivity. DCM is the predominant analysis framework for characterizing effective connectivity within distributed neuronal responses (Friston et al., 2019; Zeidman, Jafarian, Corbin, et al., 2019; Zeidman, Jafarian, Seghier, et al., 2019). Classical deterministic bilinear DCM uses a bilinear state equation with three components, including experimental (driving) inputs perturbing brain states (i.e., in our case, different velocity tactile stimuli), intrinsic connectivity in the absence of experimental perturbations, and modulations of the intrinsic connectivity induced by experimentally manipulated inputs (i.e., changes in regional couplings by tactile stimuli), which provided information concerning how much activation in source regions receiving direct inputs caused an increase/decrease in activation in target regions per unit of time. DCM is well suited to examine the directed causal relationships within the sensorimotor network during passive orofacial pneumotactile stimulation in this study.

The present study aimed to identify effective connectivity in 20 neurotypical adults’ fMRI data using DCM during orofacial pneumotactile stimuli through a 5-channel array at three saltatory velocities (5, 25, and 65 cm/s). This work is an extension of our previous studies (Custead et al., 2017; Wang et al., 2020) and will provide new information on the causal relationships between brain regions within the sensorimotor systems responsible for encoding the velocity tactile stimulation. We aimed to address the following questions on 1) whether orofacial pneumotactile stimuli (5, 25, and 65 cm/s) are processed serially from the contralateral SI to the contralateral SII or parallelly to both contralateral SI and SII, 2) how orofacial pneumotactile stimuli (5, 25, and 65 cm/s) influence interhemispheric connections between the contralateral SII and the ipsilateral SII, 3) how passive orofacial pneumotactile stimuli (5, 25, and 65 cm/s) influence the contralateral M1, and 4) what is the role of the right lobule VI in the sensorimotor networks during orofacial pneumotactile stimuli (5, 25, and 65 cm/s). Our results will provide direct evidence that passive orofacial pneumotactile stimuli with different velocities can induce functional changes in the sensorimotor network and enhance sensorimotor abilities after brain injury (i.e., due to stroke, traumatic brain injury, etc.).

## 2. METHODS

This study used a dataset that has been described in previous publications (Custead et al., 2017; Wang et al., 2020), which provides additional details regarding participants, paradigms, and fMRI data preprocessing.

### 2.1 Participants

Twenty healthy adults (mean age of 22.3, 15 females) were all right-handed, native English speakers and signed written informed consent forms to be enrolled in this study. They reported no history of neurological or psychiatric disorders. The present study was approved by the Institutional Review Board at the University of Nebraska–Lincoln (UNL).

### 2.2 Paradigms

The block-design fMRI paradigm presented each condition in a block of a 20-s task period followed by a 20-s rest period (Custead et al., 2017; Wang et al., 2020). The five task conditions consisted of 5 cm/s, 25 cm/s, 65 cm/s, “All on,” and “All off” were randomly presented. The different velocities represented the different saltation speeds of the 60-millisecond air pressure pulses through the facial array. During the task condition, the participant passively received pneumotactile stimuli to the right facial skin by the Galileo Somatosensory™ system (a multichannel pneumatic amplifier and tactile array, Epic Medical Concepts & Innovations, Inc., Mission, Kansas, K.S., U.S.A.). During the rest condition, a visual countdown on the screen was used to maintain the participant’s vigilance using E-Prime 2.0 (Psychology Software Tools, Pittsburgh, P.A., U.S.A.). Participants were instructed to pay attention to the number shown on the screen for 0.5 s to minimize brain activation in the primary visual cortex. A declining numeric countdown from 20 to 1 was used to indicate the rest period’s remaining time. To reduce the effect of fatigue, we did three runs separately and offered optional breaks between runs. Each run consisted of 20 blocks, including four blocks of 5 cm/s, four blocks of 25 cm/s, four blocks of 65 cm/s, four blocks of “All on,” and four blocks of “All off” (Custead et al., 2017; Wang et al., 2020). In total, each condition block lasted 960 s with 480-s condition segment and 480-s rest segment. Nineteen participants completed all three runs, and one participant completed two runs.

### 2.3 Image acquisition

All images were collected using a 3T Siemens Skyra MRI system (Siemens Medical Solutions, Erlangen, Germany) with a 32-channel head coil at the Center for Brain, Biology and Behavior at UNL. A high-resolution T1-weighted three-dimensional anatomical scan was acquired using magnetization-prepared rapid gradient-echo sequences (MPRAGE) with the following parameters: TR/TE/TA = 2.4 s/3.37 ms/5:35 min, flip angle = 7°, field of view = 256×256mm, spatial resolution = 1×1×1mm^3^, number of slices = 192. Following the anatomical scan, the functional MRI (fMRI) scans were recorded using a T2*-weighted echo-planar imaging (EPI) sequence with the following parameters: TR/TE/TA = 2.5 s/30 ms/800 s, voxel size = 2.5 × 2.5 × 2.5 mm^3^, flip angle = 83°, number of slices = 41, number of volumes = 320.

### 2.4 Preprocessing and general linear model

All image data from each run were concatenated and preprocessed using the SPM12 toolbox (Wellcome Department of Imaging Neuroscience, University College London, U.K.), including motion correction through spatial realignment, structural segmentation and normalization, coregistration between functional scans and anatomical scan, and smoothing with 8 mm full-width at half-maximum (FWHM) using a Gaussian Kernel.

At the first (individual) level, the general linear model (GLM) estimated the parameters for each task condition when controlling motion using six rigid-body parameters as nuisance regressors. The overall main effect of velocity (5, 25, 65 cm/s) was computed with *F*-test using analysis of variance (ANOVA). Three *T*-test contrasts included 5cm/s > rest, 25 cm/s > rest, 65 cm/s > rest. At the second (group) level, individual SPM results were pooled together for each contrast using a random-effect one-sample *T*-test. The bspmview toolbox (https://www.bobspunt.com/software/bspmview/) was used to create an axial view of slice montage. (*q* < 0.05, False discovery rate (FDR) corrected) (Benjamini & Hochberg, 1995).

### 2.5 Regions of interest and time-series extraction

The regions of interest (ROIs) were selected based on the group results for each velocity stimulus. For 5 cm/s and 25 cm/s, ROIs included bilateral primary and secondary somatosensory cortices (left SI–LSI, SII–LSII, right SI–RSI, SII–RSII), left primary motor cortex (LM1), and right cerebellar lobule VI (RVI). For 65 cm/s, ROIs included the left primary and secondary somatosensory cortices. The coordinates of each ROI from the group results were used as the center of 8 mm radius spherical volumes to search a local maximum for each individual. Contrasts for the effect of each condition (5, 25, 65 cm/s) were used to identify peak voxels. Not all participants had significantly active voxels within each ROI. For each contrast of interest, mean-corrected (by an F-contrast for the effects of velocity) time series from each participant were extracted within 8 mm radius spherical volumes centered on each ROI using the first eigenvariate of voxels above a threshold of *p* < 0.001 (uncorrected).

### 2.6 Dynamic causal modeling

DCM was used in the present study to test hypotheses about the neuronal mechanisms that underlie experimental measurements of brain responses (Stephan et al., 2010). DCM uses bilinear differential equations with three components, including experimental (driving) inputs perturbing brain states (i.e., for our study, different velocity tactile stimuli elicit the cortex) (DCM.C matrix), intrinsic connectivity in the absence of experimental perturbations (DCM.A matrix), and changes (modulations) of the intrinsic connectivity induced by experimentally manipulated inputs (i.e., for our study, changes in regional couplings by tactile stimuli, which provided information concerning how much activation in source regions receiving direct inputs caused an increase/decrease in activation in target regions per unit of time) (DCM.B matrix) (Stephan, Weiskopf, Drysdale, Robinson, & Friston, 2007). DCM can infer causal mechanisms in the brain networks and how external stimuli can change the causal relationships in the networks (Stephan et al., 2010; Stephan et al., 2007). The model space in DCM analysis is constructed by hypotheses about the effective connectivity in the brain networks of interest.

To test whether pneumotactile saltatory stimuli to the right facial skin are processed using serial mode or parallel mode for each velocity, we constructed a set of six models for 5 and 25 cm/s and a set of two models for 65 cm/s (see Figure 1). We included the intrinsic reciprocal connectivity (all forward and backward fixed connections) between the ROIs for each task condition. The pneumotactile stimuli were the driving inputs perturbing brain states either through the left SI only or both left SI and SII. To test whether velocity encoding modulates interhemispheric connection between the contralateral left SII and the ipsilateral right SII in forward-only mode, backward-only mode, or both forward and backward modes for 5 and 25 cm/s. We also included modulations of the intrinsic connectivity induced by either 5 or 25 cm/s stimuli. To examine how different velocities modulate the intrinsic connections between the left SI, SII, and M1, we constructed a set of three models for 5 and 25 cm/s (see Figure 2), including modulating through forward connection from the left SI to left M1, or from the left SII to left M1, or from both the left SI and SII to left M1. To examine the cerebellum’s role in somatosensory networks, we constructed a set of nine models, including the intrinsic reciprocal connectivity between the ROIs for 5 and 25 cm/s (see Figure 3). The external velocity stimuli modulate the right cerebellar lobule VI through forward, or backward, or both connections to other ROIs (the left SI, SII, and M1).

**Figure 1.**
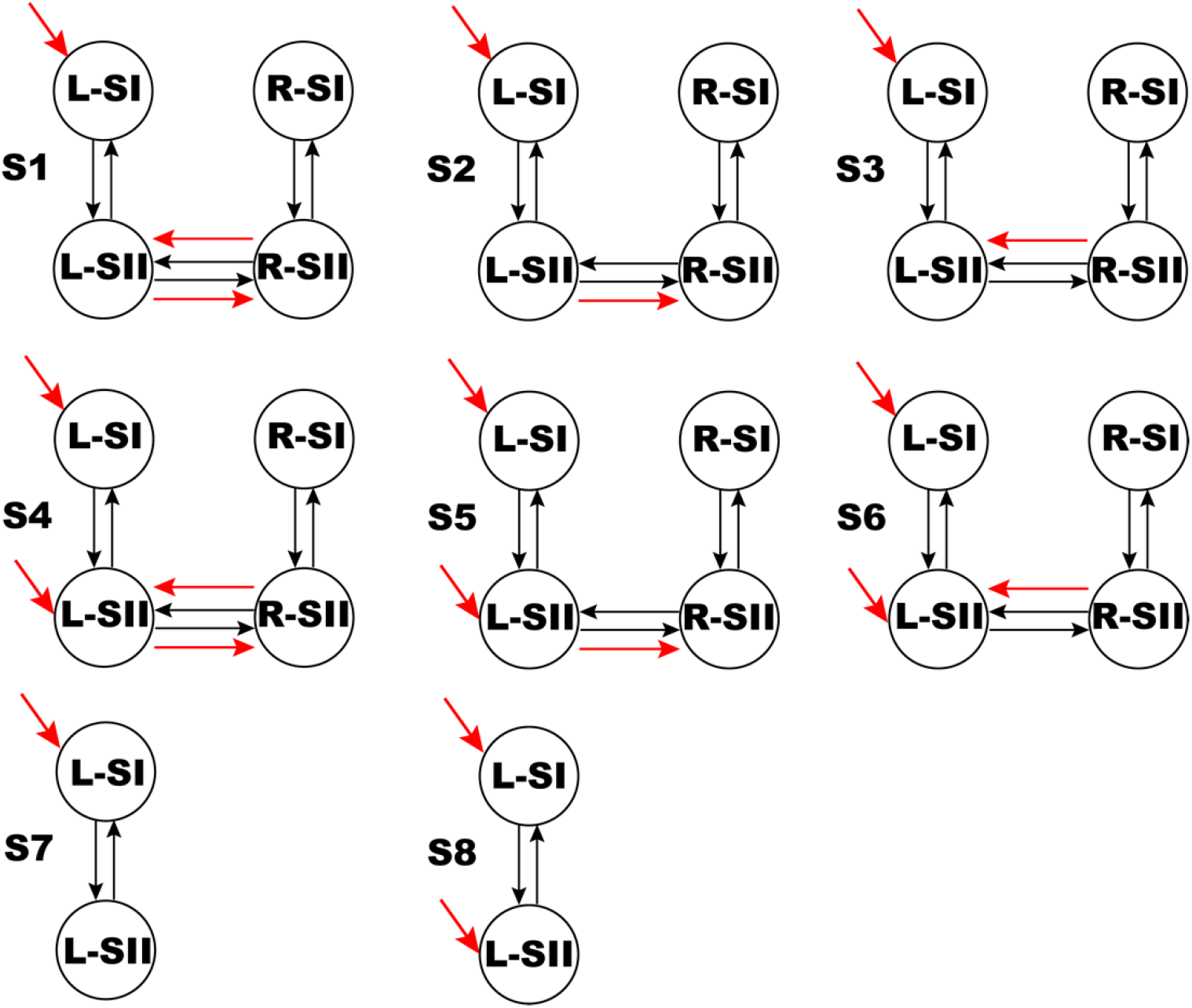
L-SI: left primary somatosensory cortex, L-SII: left secondary somatosensory cortex, R-SI: right primary somatosensory cortex, R-SII: right secondary somatosensory cortex. Black arrows represent the intrinsic connections, Red arrows represent the modulatory connectivity and driving inputs to L-SI and L-SII. Models S1–S6 were examined for 5 and 25 cm/s. Models S7– S8 were examined for 65 cm/s.

**Figure 2.**
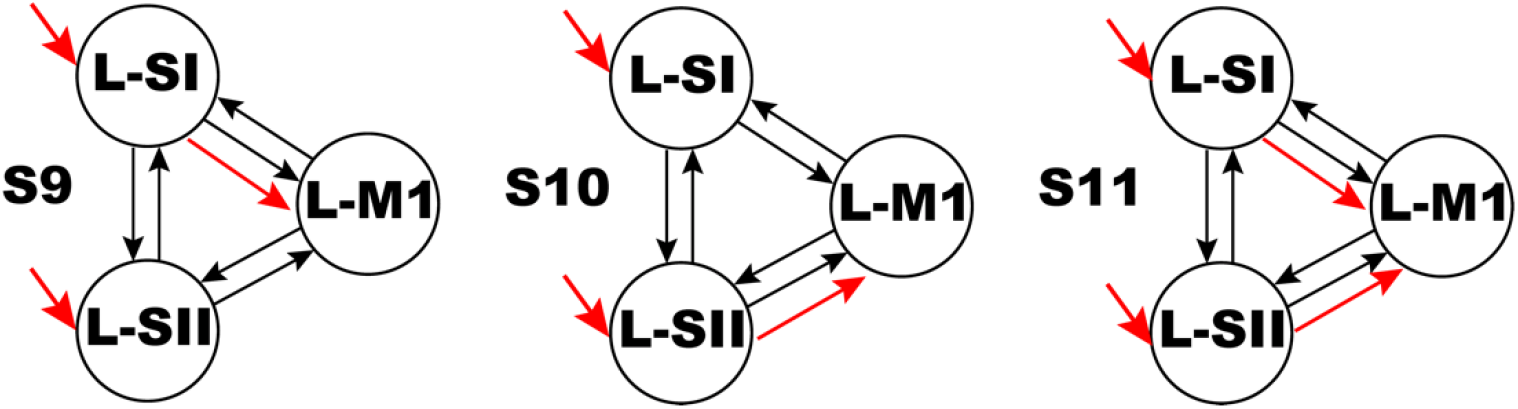
L-SI: left primary somatosensory cortex, L-SII: left secondary somatosensory cortex, R-SI: right primary somatosensory cortex, R-SII: right secondary somatosensory cortex, L-M1: left primary motor cortex. Black arrows represent the intrinsic connections, Red arrows represent the modulatory connectivity and driving inputs to L-SI and L-SII. Models S9–S11 were examined for 5 and 25 cm/s.

**Figure 3.**
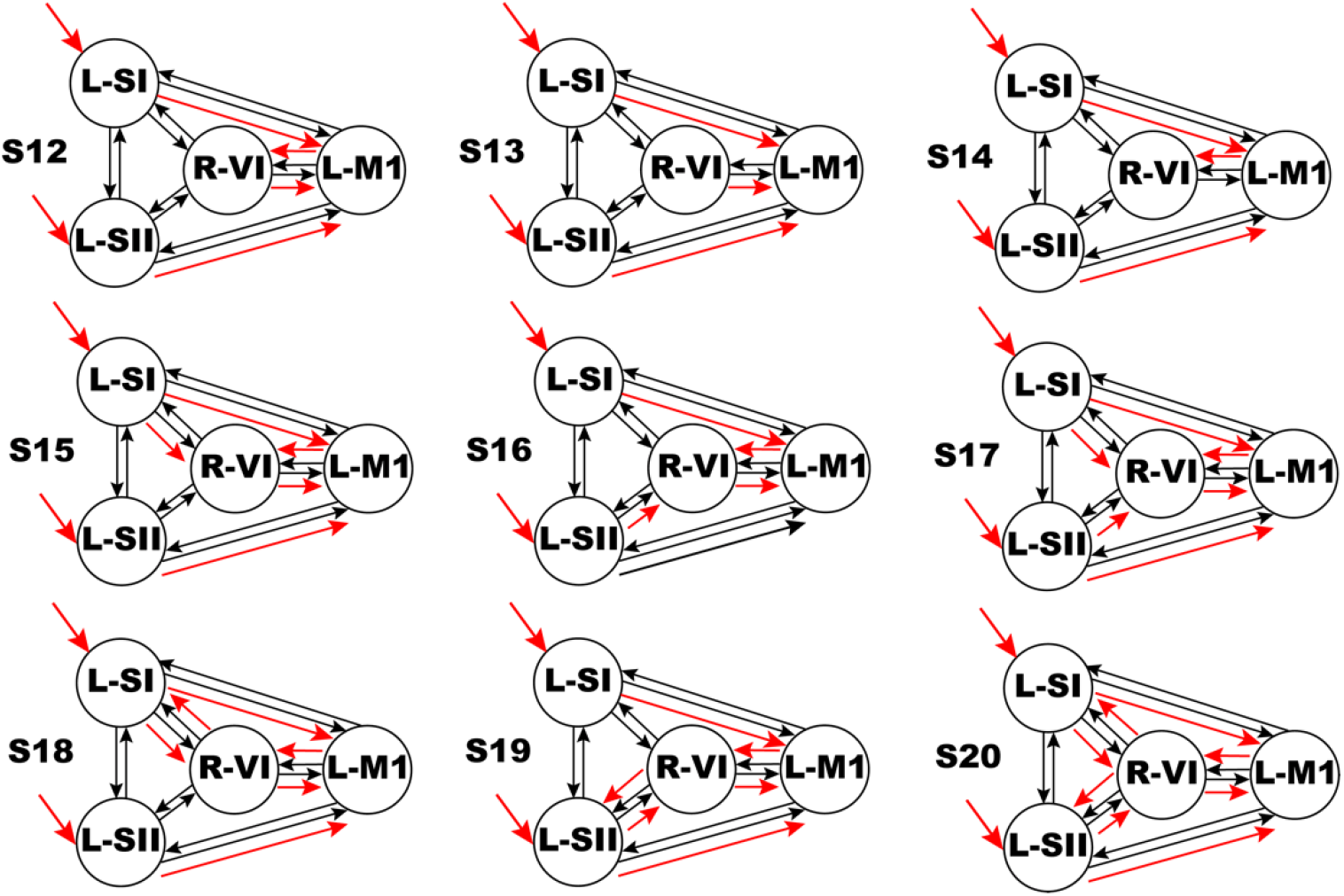
L-SI: left primary somatosensory cortex, L-SII: left secondary somatosensory cortex, R-SI: right primary somatosensory cortex, R-SII: right secondary somatosensory cortex, L-M1: left primary motor cortex, R-VI: right cerebellar lobule VI. Black arrows represent the intrinsic connections, Red arrows represent the modulatory connectivity and driving inputs to L-SI and L-SII. Models S12–S20 were examined for 5 and 25 cm/s.

### 2.7 Bayesian model selection and inference on parameters

The Bayesian model selection (BMS) compares each model’s log evidence approximated using the free energy to select the winning model that explains the data as accurately as possible and has minimal complexity (Stephan et al., 2010; Stephan et al., 2007). The model with the highest posterior probability is also the model with the most substantial evidence (Kass & Raftery, 1995; Rosa, Friston, & Penny, 2012). Bayes factors have been used to compare two models (Kass & Raftery, 1995). Under the uniform priors and Bayes’ rule, a posterior model probability greater than 95% is equivalent to a Bayes factor greater than 20, which provides strong evidence in favor of one model over the other (Kass & Raftery, 1995; Rosa et al., 2012). A Bayes factor of 1 to 3 corresponding to a posterior model probability of 50-75% indicates weak evidence. A Bayes factor of 3 to 20 corresponding to a posterior model probability of 75-95% suggests positive evidence. We used a random-effects (RFX) approach for model selection at the group level, which allows each participant to have a different best model and computes the probability of all participants’ data given each model. The model with high exceedance probability (EP) is the winning model. For the winning model, we considered the winning model’s parameters as random effects in the population (i.e., velocity stimuli induced changes in connection strengths) (Stephan et al., 2010). Thus, the Bayesian model averaging (BMA) approach was used to compute the weighted averages of each model parameter. The weighting is determined by the posterior probability of each model (Stephan et al., 2010). Furthermore, the BMA values of the winning model were reported for each DCM analysis. We marked the connectivity parameters with the probability that the posterior estimate of the parameter is not zero greater than 85%. In addition, the modulation effects of velocity stimuli were compared using BMA results with one-sample paired *t*-tests with FDR correction for multiple comparisons (Benjamini & Hochberg, 1995) (*q* < 0.05, FDR corrected).

## 3. RESULTS

### 3.1 GLM random-effects analysis

The group results were shown in Figure 4, as identified in our previous report of this dataset (Custead et al., 2017; Wang et al., 2020). In the current DCM analysis, we focused on six ROIs for 5 and 25 cm/s and two ROIs for 65 cm/s. The group averaged coordinates were reported in Table 1.

**Table 1.** Group averaged coordinates of ROIs’ center in MNI space.

| ROI | 5 cm/s |  |  | 25 cm/s |  |  | 65 cm/s |  |  |
| --- | --- | --- | --- | --- | --- | --- | --- | --- | --- |
|  | X, Y, Z | N | T | X, Y, Z | N | T | X, Y, Z | N | T |
| Left SI | -56, -22, 45 | 19 | 6.3 | -51, -20, 45 | 20 | 7.8 | -53, -21, 50 | 17 | 7.8 |
| Left SII | -54, -28, 24 | 17 | 5.7 | -54, -25, 23 | 19 | 5.4 | -53, -22, 26 | 15 | 4.5 |
| Right SI | 58, -18, 42 | 13 | 3.8 | 58, -15, 37 | 13 | 3.4 |  |  |  |
| Right SII | 58, -20, 27 | 13 | 4.0 | 56, -22, 24 | 13 | 4.0 |  |  |  |
| Left M1 | -47, -23, 51 | 17 | 3.6 | -46, -23, 52 | 17 | 4.3 |  |  |  |
| Right VI | 26, -57, -24 | 7 | 3.6 | 26, -56, -23 | 5 | 3.4 |  |  |  |

N: the number of participants who had significantly active voxels within the ROI, T: averaged T-values, SI: primary somatosensory cortex, SII: secondary somatosensory cortex, M1: primary motor cortex. Right VI: Right cerebellar lobule VI.

**Figure 4.**
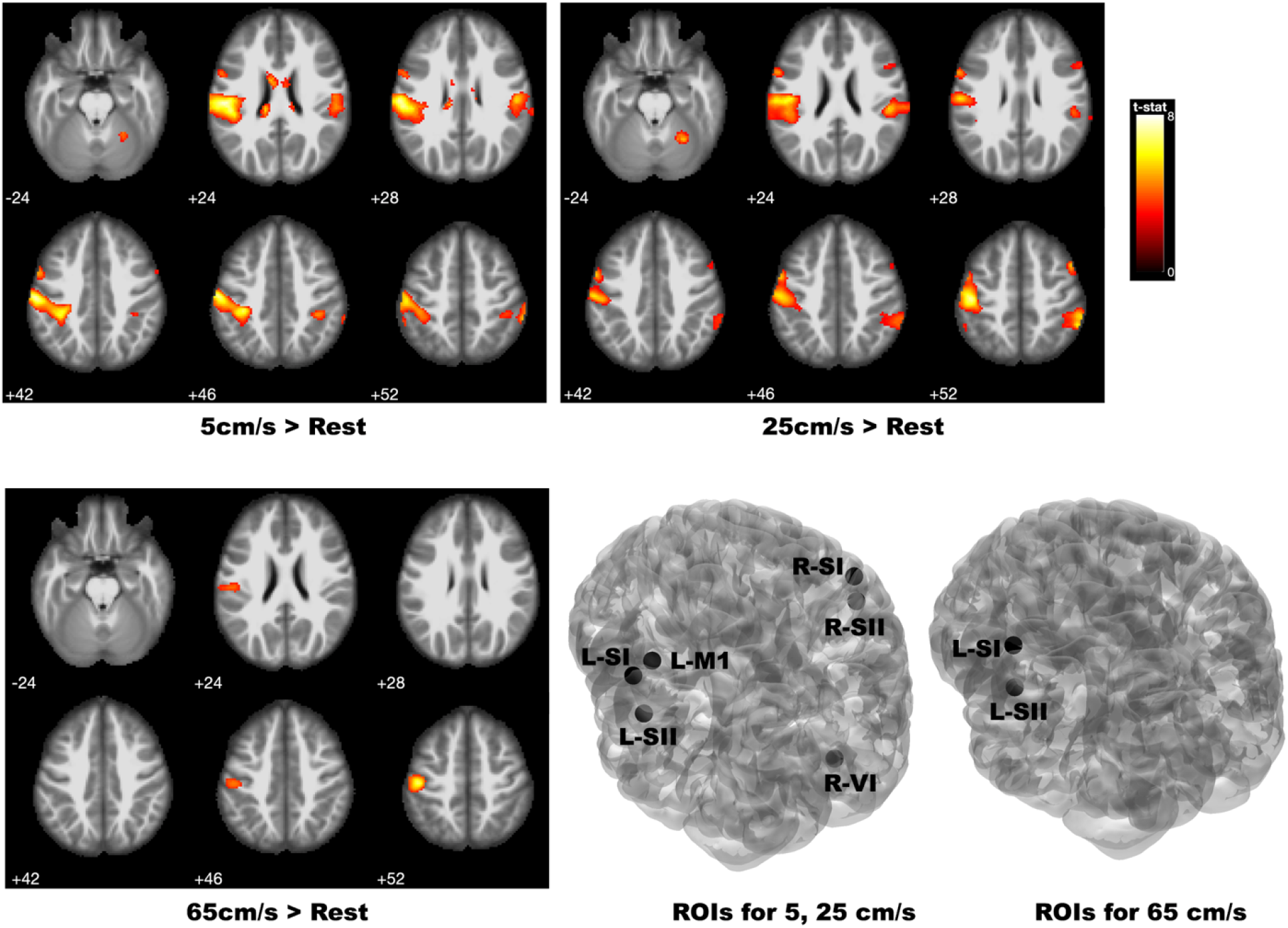
Group results for 5cm/s > Rest, 25 cm/s > Rest, 65 cm/s > Rest. ROIs for 5, 25, 65 cm/s. L-SI: left primary somatosensory cortex, L-SII: left secondary somatosensory cortex, R-SI: right primary somatosensory cortex, R-SII: right secondary somatosensory cortex, L-M1: left primary motor cortex, R-VI: right cerebellar lobule VI. Images are in neurological view: the left side of the image corresponds to the left side of the brain, and the right side of the image corresponds to the right side of the brain.

### 3.2 DCM model comparison and selection

#### 3.2.1 Driving inputs and interhemispheric connections

For both 5 and 25 cm/s, the winning model was S5 with the highest exceedance probability (5 cm/s: 85%, 25 cm/s: 51%) (Figure 5). Somatosensory stimuli were driving inputs to both left SI and SII, and sensory stimuli only modulated the forward connection from contralateral left SII to the ipsilateral right SII. Based on Bayes’ rule, there was strong evidence that the driving inputs entered the sensorimotor network through both left SI and SII for 5 cm/s (i.e., an exceedance probability of 85% relative to 0.7% for the best model among S1 to S3 with the driving inputs to the left SI only). For 25 cm/s, S5 with the highest exceedance probability (51%) was the winning model. However, there was weak evidence in favor of S5 over S3 (27%) since the Bayes factor was 1.5 computed by the ratio of model posterior means (0.3/0.2 = 1.5). For 65 cm/s, the winning model was S8 with the highest exceedance probability (55%) (Figure 5). A Bayes factor of 1 suggested that there was weak evidence in favor of S8 over S7 (45%). The driving inputs to both contralateral left SI and SII provided strong evidence on a parallel mode of processing for 5 cm/s and weak evidence on a parallel mode of processing for 25 and 65 cm/s. In addition, the 5 and 25 cm/s velocity stimuli both modulated the inter-hemispheric connection through the forward connection from the contralateral left SII to the ipsilateral right SII.

**Figure 5.**
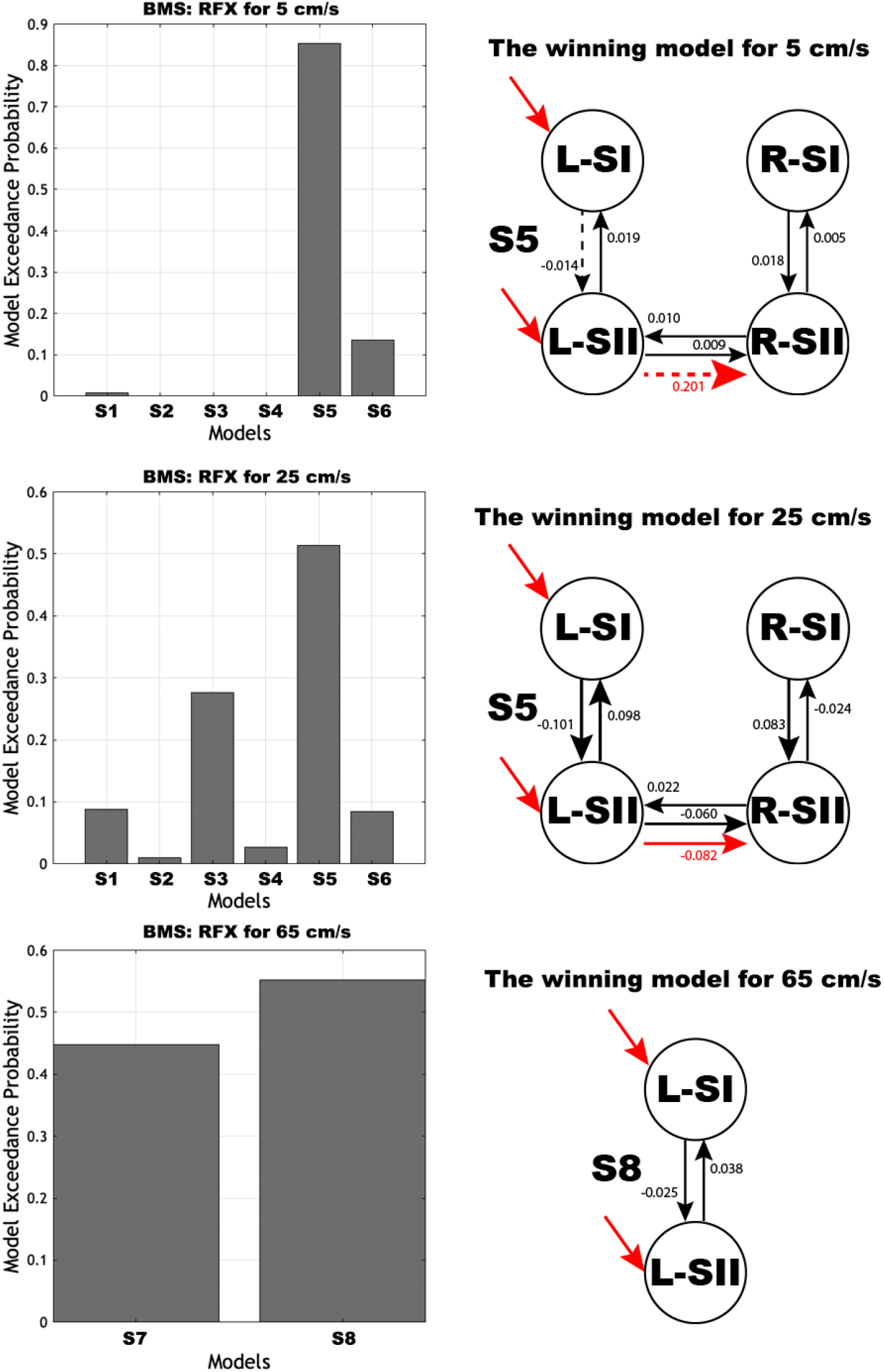
BMS: RFX results for each velocity and the winning model for each velocity. The dash line indicates probability that the posterior estimate of the parameter is not zero is less than 85%. L-SI: left primary somatosensory cortex, L-SII: left secondary somatosensory cortex, R-SI: right primary somatosensory cortex, R-SII: right secondary somatosensory cortex. Black arrows represent the intrinsic connections, Red arrows represent the modulatory connectivity and driving inputs to L-SI and L-SII. The line’s thickness is determined by the connectivity parameter.

#### 3.2.2 Cross-modality Plasticity

For 5 cm/s, the winning model was S10 with the highest exceedance probability of 50% (Figure 6). The 5 cm/s velocity stimuli modulated only forward connection from the left SII to the left M1. However, there was weak evidence in favor of S10 over S11 (49%) since the Bayes factor was one, computed by the ratio of model posterior means (0.4/0.4 = 1). For 25 cm/s, the winning model was S11 with the highest exceedance probability of 34% (Figure 6). The 25 cm/s velocity stimuli modulated both forward connections from the left SI to the left M1 and the left SII to the left M1. But there was weak evidence in favor of S11 over S9 (Bayes factor = 1) or S10 (Bayes factor = 1).

**Figure 6.**
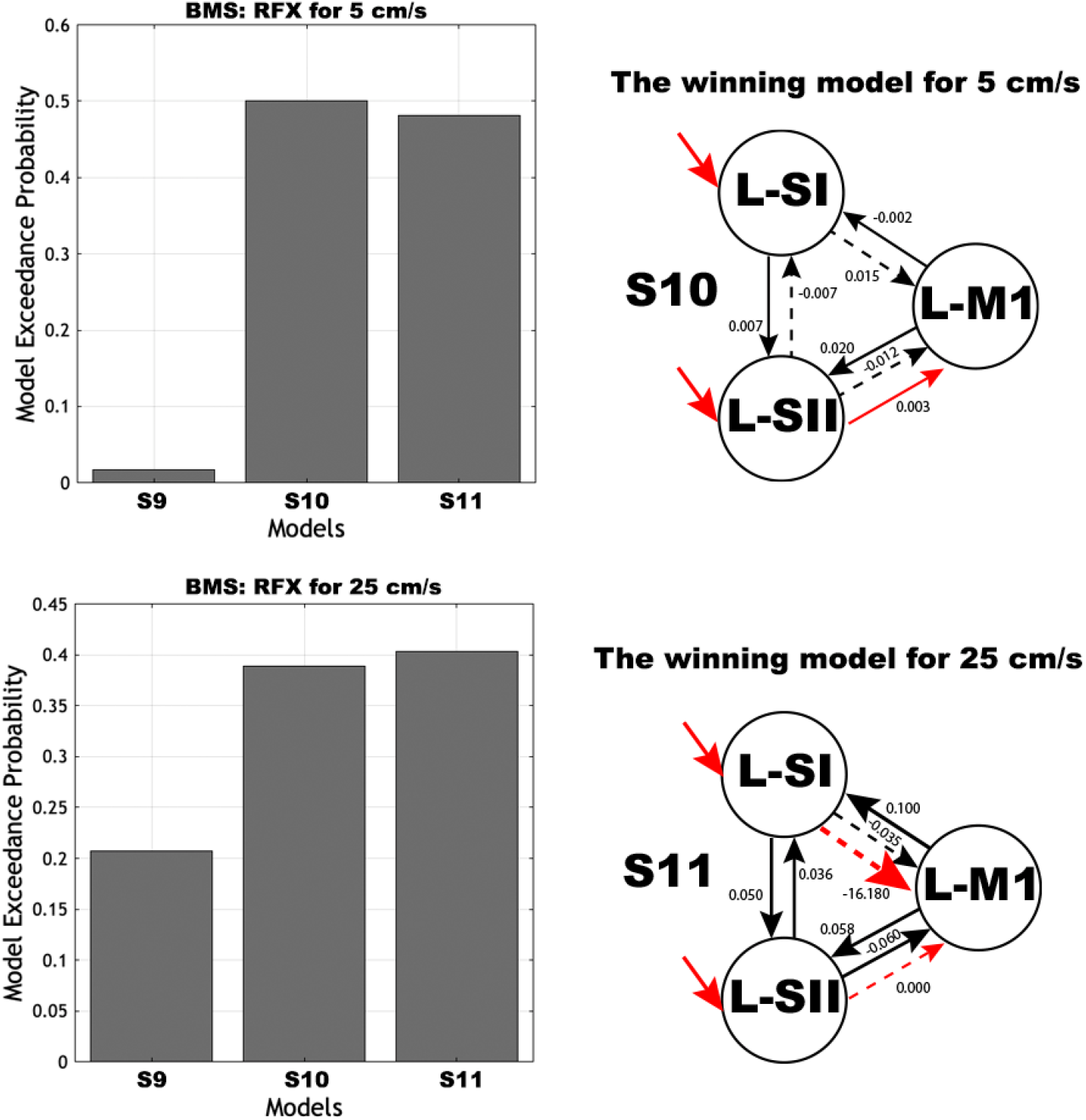
BMS: RFX results for each velocity and the winning model for each velocity. The dash line indicates probability that the posterior estimate of the parameter is not zero is less than 85%. L-SI: left primary somatosensory cortex, L-SII: left secondary somatosensory cortex, R-SI: right primary somatosensory cortex, R-SII: right secondary somatosensory cortex, L-M1: left primary motor cortex. Black arrows represent the intrinsic connections, Red arrows represent the modulatory connectivity and driving inputs to L-SI and L-SII. The line’s thickness is determined by the connectivity parameter.

#### 3.2.3 Feedforward and feedback loops of cortico-cerebral networks

For 5 cm/s, the winning model was S20 with the highest exceedance probability of 36% (Figure 7). The 5 cm/s velocity stimuli modulated forward and backward connections between the right VI to the left SI, SII, and M1. However, there was weak evidence in favor of S20 over the second-best model S18 with the exceedance probability of 21% since the Bayes factor was 1.3. For 25 cm/s, the winning model was S18 with the highest exceedance probability of 41% (Figure 7). The 25 cm/s velocity stimuli modulated forward and backward connections between the right VI to the left SI and M1. There was weak evidence in favor of S18 over the second-best model S18 with the exceedance probability of 18% since the Bayes factor was 1.5.

**Figure 7.**
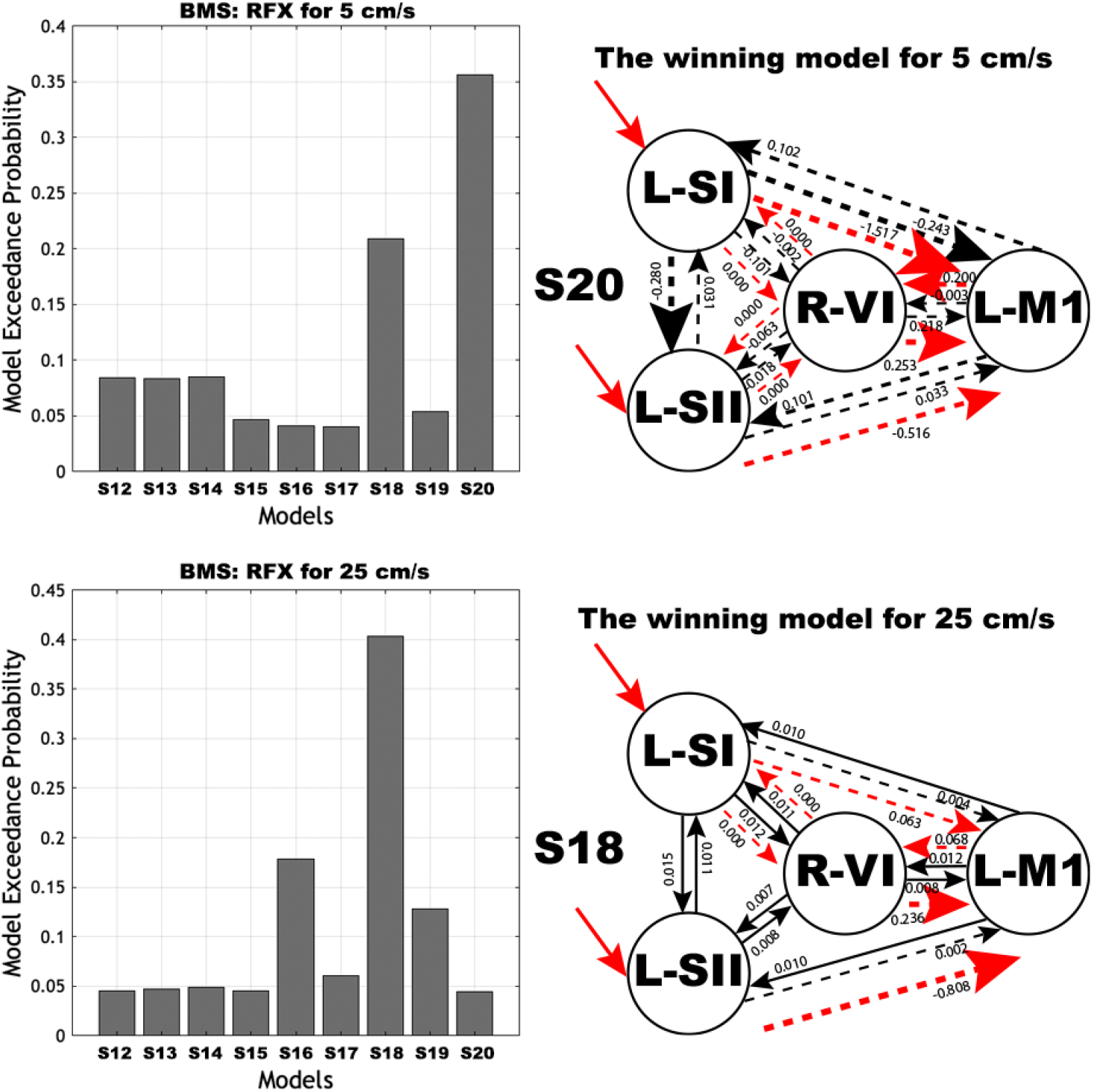
BMS: RFX results for each velocity and the winning model for each velocity. The dash line indicates probability that the posterior estimate of the parameter is not zero is less than 85%. L-SI: left primary somatosensory cortex, L-SII: left secondary somatosensory cortex, R-SI: right primary somatosensory cortex, R-SII: right secondary somatosensory cortex, L-M1: left primary motor cortex, R-VI: right cerebellar lobule VI. Black arrows represent the intrinsic connections, Red arrows represent the modulatory connectivity and driving inputs to L-SI and L-SII. The line’s thickness is determined by the connectivity parameter.

### 3.3 Connectivity parameters of the winning models

#### 3.3.1 Driving inputs and interhemispheric connections

BMA results of connectivity parameters for S5 and S8 models were shown in Table 2 and Figure 5. Positive numbers indicate excitation, and negative numbers indicate inhibition. For all velocities, the driving inputs to the left SI and SII researched significance. There was no significant difference between the modulation effect of 5 cm/s and the modulation effect of 25 cm/s on forward (*p* = 0.096) connection from the left SII to the right SII.

**Table 2.** BMA results of connectivity parameters of S5 and S8.

| Connections | Intrinsic strength 5 cm/s (Hz) | Effect of 5 cm/s (no unit) | Intrinsic strength 25 cm/s (Hz) | Effect of 25 cm/s (no unit) | Intrinsic strength 65 cm/s (Hz) |
| --- | --- | --- | --- | --- | --- |
|  | S5 model |  | S5 model |  | S8 model |
| L-SI → L-SII | -0.014 | n/a | -0.101† | n/a | -0.025† |
| L-SI ← L-SII | 0.019† | | 0.098† | | 0.038† |
| L-SII → R-SII | 0.009† | 0.201 | -0.060† | -0.082† | n/a |
| L-SII ← R-SII | 0.010† | n/a | 0.022† | n/a | |
| R-SI → R-SII | 0.018† | n/a | 0.083† | n/a | |
| R-SI ← R-SII | 0.005† | | -0.024† | | |

Note: L-SI: left primary somatosensory cortex, L-SII: left secondary somatosensory cortex, R-SI: right primary somatosensory cortex, R-SII: right secondary somatosensory cortex, L-M1: left primary motor cortex. † Probability that the posterior estimate of the parameter is not zero is greater than 85%. n/a: not applicable.

#### 3.3.2 Cross-modality Plasticity and Feedforward and feedback loops of cortico-cerebral networks

BMA results of connectivity parameters for S10, S11, S20, and S18 models were shown in Table 3 and Figure 6, 7. Positive numbers indicate excitation and negative numbers indicate inhibition. There was no significant difference between the modulation effect of 5 cm/s and the modulation effect of 25 cm/s on forward connection (*p* = 0.331) from the left SI to the left M1 or forward connection (*p* = 0.117) from the left SII to the left M1. There was no significant difference between the modulation effect of 5 cm/s and the modulation effect of 25 cm/s on the forward connections from the left SI to the left M1 (*p* = 0.365) and from the left SII to the left M1 (*p* = 0.408). In addition, there was no significant difference between the modulation effect of 5 cm/s and the modulation effect of 25 cm/s on the forward connections from the left SI to the right VI (*p* = 0.788), from the left SII to the right VI (*p* = 0.219), and from the left M1 to the right VI (*p* = 0.573), as well as the backward connections from the right VI to the left SI (*p* = 0.189), from the right VI to the left SII (*p* = 0.177), and the right VI to the left M1 (*p* = 0.534).

**Table 3.** BMA results of connectivity parameters of S10, S11, S18, and S20.

| Connections | Intrinsic strength<br>5 cm/s | Effect<br>of 5<br>cm/s | Intrinsic strength<br>25 cm/s | Effect<br>of 25<br>cm/s | Intrinsic strength<br>5 cm/s | Effect<br>of 5<br>cm/s | Intrinsic strength<br>25 cm/s | Effect<br>of 25<br>cm/s |
| --- | --- | --- | --- | --- | --- | --- | --- | --- |
|  | S10 model |  | S11 model |  | S20 model |  | S18 model |  |
| L-SI → L-SII | 0.007† | n/a | 0.050† | n/a | -0.280 | n/a | 0.015† | n/a |
| L-SI ← L-SII | -0.007 | | 0.036† | | 0.031 | | 0.011† | |
| L-SI → L-M1 | -0.002 | | -0.035 | -16.180 | -0.243 | -1.517 | 0.004 | 0.063 |
| L-SI ← L-M1 | 0.015† | | 0.100† | n/a | 0.102 | n/a | 0.010† | n/a |
| L-SII → L-M1 | -0.012 | 0.003† | -0.060† | 0 | 0.033 | -0.156 | 0.002 | -0.808 |
| L-SII ← L-M1 | 0.020† | n/a | 0.058† | n/a | 0.101 | n/a | 0.010† | n/a |
| L-SI → R-VI | n/a | | | | -0.002 | 0 | 0.011† | 0 |
| L-SI ← R-VI | | | | | -0.101 | 0 | 0.012† | 0 |
| L-SII → R-VI | | | | | -0.018 | 0 | 0.008† | n/a |
| L-SII ← R-VI | | | | | -0.063 | 0 | 0.007† | |
| L-M1 → R-VI | | | | | -0.003 | 0.200 | 0.012† | 0.068 |
| L-M1 ← R-VI | | | | | 0.218 | 0.253 | 0.008† | 0.236 |

Note: L-SI: left primary somatosensory cortex, L-SII: left secondary somatosensory cortex, R-SI: right primary somatosensory cortex, R-SII: right secondary somatosensory cortex, L-M1: left primary motor cortex, R-VI: right cerebellar lobule VI. † Probability that the posterior estimate of the parameter is not zero is greater than 85%. n/a: not applicable.

## 4. DISCUSSION

Our previous functional connectivity study identified similarities of neuronal networks in the contralateral brain regions and differences of neuronal networks in the ipsilateral brain regions among three-velocity pneumotactile stimuli (5, 25, 65 cm/s) (Wang et al., 2020). Our previous study was unable to identify the causal relationships among brain regions under different pneumotactile stimuli. Thus, the present study aimed to use DCM to examine velocity pneumotactile stimulus-related influences between brain regions in the sensorimotor networks during passive saltatory pneumotactile stimuli (5, 25, 65 cm/s), which has not been reported previously. First, the winning model S5 for 5 and 25 cm/s and S8 for 65 cm/s defined the contralateral left SI and SII as the most likely sources of the driving inputs within the sensorimotor network. In our model space, there were two possible ways for pneumotactile stimuli to enter the sensorimotor network. Our results provided strong evidence that the cortical network supporting higher-order processing of the facial pneumotactile stimuli (5 cm/s) involved parallel processing with driving inputs to both contralateral SI and SII. For medium and high velocities (25 and 65 cm/s), the most likely sources for the driving inputs were both left SI and SII, but the evidence was relatively weak. Second, model S5 provides direct evidence for the modulation effect on the forward interhemispheric connection from the contralateral left SII to the right SII, especially at the 25 cm/s pneumotactile stimuli. Third, the S10 and S11 models provided direct evidence that the velocity pneumotactile stimuli influenced the left M1 through the left SI and SII. Our results indicate that passive pneumotactile stimulation may positively impact motor function rehabilitation. Furthermore, the S18 and S20 models provided direct evidence that the right cerebellar lobule VI plays a role in the sensorimotor network through forward and backward neuronal pathways. The slow velocity of 5 cm/s pneumotactile stimuli modulated both forward and backward connections between the right cerebellar lobule VI and the contralateral left SI, SII, and M1. The medium velocity of 25 cm/s pneumotactile stimuli modulated both forward and backward connections between the right cerebellar lobule VI and the contralateral left SI and M1, whereas the 65 cm/s pneumotactile stimuli did not elicit activation in the right cerebellar lobule VI in the majority of participants.

### 4.1 Parallel processing of velocity pneumotactile stimuli

In the present study, the right lower face was passively and noninvasively stimulated with air pressure pules from a spatial array of TAC-Cells (Wang et al., 2020). The pneumotactile stimuli are received through the cutaneous mechanoreceptors in the facial skin to the brain stem and then to the thalamus. Our DCM results provide strong evidence for a network able to parallel process of low velocity (5 cm/s) orofacial pneumotactile stimuli, which is in agreement with some studies (Klingner et al., 2015; Liang et al., 2011; Song et al., 2021) and also contrasts other findings (Disbrow et al., 2001; Kalberlah et al., 2013; Khoshnejad et al., 2014). Liang et al. has reported that non-nociceptive and nociceptive somatosensory inputs (electrical pulses to the right ankle that activate all subpopulation of fast-conducting myelinated Aß fibers) are processed in parallel from the thalamus to S1 and from the thalamus to S2 using DCM of fMRI data (Liang et al., 2011). Another MEG study (Klingner et al., 2015) used electrical median nerve stimulus to the right wrist and found a parallel processing pathway to both contralateral SI and SII using DCM. A recent fMRI study (Song et al., 2021) used both nociceptive laser stimuli and electrical tactile stimuli to the right foot and identified parallel ascending pathways for both types of sensory stimuli through the thalamus to both SI and SII using DCM. The present study used pneumotactile stimuli to the right lower face area and found parallel processing of orofacial pneumotactile stimuli using DCM, which has not been reported previously. Contrary to our findings, two DCM studies supported serial processing from the contralateral SI to SII in response to innocuous and noxious electrical stimuli to the right sural nerve (Khoshnejad et al., 2014), as well as tactile vibratory stimuli to the left middle and index fingers (Kalberlah et al., 2013). The contradicting findings could result from various experimental settings (i.e., electrical versus tactile vibratory stimuli, stimulation to fingers/medial nerve versus foot/ankle). Additionally, the present study provides weak evidence in favor of parallel processing over serial processing of the orofacial pneumotactile stimuli for medium (25 cm/s) and high (65 cm/s) velocity. There was also significant intrinsic connectivity from the left SI to SII for 25 and 65 cm/s velocities. These findings from the present study suggested the coexistence of the parallel and serial processing theories regarding medium-to high-velocity pneumotactile processing, consistent with other studies (Chung et al., 2014; Cruccu et al., 2008). Low velocity (5 cm/s) orofacial pneumotactile stimuli result in parallel processing, whereas medium (25 cm/s) and high (65 cm/s) velocities recruit both serial and parallel processing types (from the present study).

### 4.2 Interhemispheric connection modulated by velocity pneumotactile stimuli

In this study, the S5 model optimally encoded the effect of velocity pneumotactile stimuli on a forward connection from the contralateral SII to the ipsilateral SII, suggesting interhemispheric modulation of effective connectivity in the SII due to the velocity pneumotactile stimuli to the lower face. Previous EEG and MEG studies have shown that unilateral somatosensory stimuli activated bilateral SII, and the activation of the ipsilateral SII was delayed (Hoechstetter et al., 2001; Stancak et al., 2002). Our DCM results are consistent with previous work and suggest that the interhemispheric modulation effect from the contralateral SII to the ipsilateral SII might be related to bidirectional transcallosal connections linking both SII. Animal studies have found relatively dense transcallosal fibers connecting the contralateral SII and ipsilateral SII (Jones & Powell, 1969a; Pandya & Vignolo, 1968; Picard et al., 1990). Moreover, we identified an excitatory forward connection from the contralateral SII to the ipsilateral SII for low velocity (5 cm/s) stimuli and inhibitory connection for medium velocity (25 cm/s) stimuli, which has not been reported previously. Compared to 5 cm/s, 25 cm/s has a higher temporal density of air-pulse stimulation and lower perception accuracy (Lamb, 1983). Thus, the medium velocity (25 cm/s) stimuli might be more affected by adaptation or repetition-suppressing process (Hollins, Delemos, & Goble, 1991; Popescu, Barlow, Venkatesan, Wang, & Popescu, 2013; Yang et al., 2014), whereas the 5-cm/s velocity stimuli might be processed as discrete stimuli instead of a constant motion across the skin (Depeault, Meftah El, & Chapman, 2013; Wacker, Spitzer, Lutzkendorf, Bernarding, & Blankenburg, 2011). The medium velocity (25 cm/s) stimuli modulated the inhibitory forward connection from the contralateral SII to the ipsilateral SII, suggesting interhemispheric inhibition within SII for encoding medium velocity pneumotactile stimuli. Similarly, the interhemispheric inhibition within SII has been related to more complex bilateral receptive fields in SII than in SI for comparing spatial features of objects (Jung et al., 2012). For the high velocity (65 cm/s) stimuli, the interhemispheric modulation effect cannot be evaluated due to the lack of activation in the ipsilateral SII. This result is in agreement with our previous study (Wang et al., 2020) and suggested the high velocity exceeded the optimal range for moving tactile stimuli, which has been reported to be 3–25 cm/s for the face (Dreyer, Duncan, Wong, & Whitsel, 1979; Edin, Essick, Trulsson, & Olsson, 1995; Whitsel et al., 1986).

Our results also showed the significant intrinsic forward and backward connections between the contralateral left SII and ipsilateral right SII for 5 and 25 cm/s velocity stimuli. The average (baseline) effective connection between the left SII and right SII suggested the existence of interhemispheric connection within the sensorimotor network. The low velocity (5 cm/s) stimuli elicited excitatory connection from the contralateral SII to the ipsilateral SII, which aligns with other studies using either pressure or electrical stimulation (Chung et al., 2014; Khoshnejad et al., 2014). Khoshnejad et al. (Khoshnejad et al., 2014) used electrical stimulation on the right sural nerve with low, moderate, and high-intensity levels and identified the excitatory connection from the contralateral SII to the ipsilateral SII for low (innocuous) and moderate (moderate-noxious) intensity levels but not for the high (high-noxious) intensity level. No participant in the present study reported discomfort or pain sensation. Therefore, pain-related neuronal networks may influence our results. However, the medium velocity (25 cm/s) stimuli elicited inhibitory connection from the contralateral SII to the ipsilateral SII. This result might be due to the complexity of the 25 cm/s velocity stimuli and effect from adaptation or repetition-suppressing process (Hollins et al., 1991; Popescu et al., 2013; Yang et al., 2014).

### 4.3 Cross-modality plasticity

Our DCM results provided weak evidence in favor of S10 over S11 for the low velocity (5 cm/s) and in favor of S11 over S10 for the medium velocity (25 cm/s). Nevertheless, both S10 and S11 indicated that the passive somatosensory inputs could modulate the primary motor cortex through either forward connection from the contralateral SII to the contralateral M1 (5 cm/s) or forward connections from both contralateral SI and SII to the contralateral M1 (25 cm/s). Therefore, our findings cannot definitely confirm which pathway passive somatosensory inputs influence the primary motor cortex. Still, our results support that the passive somatosensory inputs can affect motor function. The cross-modality plasticity theory suggested that somatosensory stimuli could evoke neural responses to promote motor learning (Ackerley, Borich, Oddo, & Ionta, 2016; Ladda et al., 2014; Ludlow et al., 2008; Nasir et al., 2013; Sanes, 2003; Sanes & Donoghue, 2000; Veldman et al., 2018). Research has shown that the orofacial sensorimotor system is essential for sucking, swallowing, and speech production (Barlow, 1998; Barlow & Bradford, 1996; Barlow & Estep, 2006; Barlow, Lund, Estep, & Kolta, 2010; Barlow & Stumm, 2010; Sessle et al., 2007; Sessle et al., 2005; Smith, 2016). Thus, our DCM results support that the passive pneumotactile stimulation could effectively modulate sensory and motor system to impact motor rehabilitation positively, in agreement with other studies (Ahn, Lee, & Hwang, 2016a; Chen et al., 2018b; Dinse & Tegenthoff, 2015; Heba et al., 2017; Macaluso, Cherubini, & Sabatini, 2007). This is an important step for developing future early neurorehabilitation protocols. For instance, individuals cannot perform active movement rehabilitation tasks after severe brain injury (i.e., stroke, traumatic brain injury, etc.) and can benefit from passive sensory stimulation paradigms. Moreover, during the critical period when neural plasticity is the highest, if the passive tactile stimuli can be used to stimulate the sensory and motor system, it may improve the motor rehabilitation’s outcomes later according to the cross-modality plasticity theory (Ackerley et al., 2016; Barlow, Custead, Lee, Hozan, & Greenwood, 2020; Blatow et al., 2011; Ladda et al., 2014; Ludlow et al., 2008; Nasir et al., 2013; Sanes, 2003; Sanes & Donoghue, 2000; Veldman et al., 2018). A recent EEG study has reported that passive unilateral somatosensory electrical stimulation can improve skill acquisition, consolidation, and interlimb transfer by increasing sensorimotor activity and connectivity (Veldman et al., 2018). Our noninvasive low velocity pulsed pneumotactile stimuli on the right lower face can modulate forward connection from the contralateral SII to the contralateral M1, supporting the cross-modality plasticity theory. Further research is needed to provide strong evidence on our findings and uncover the neural mechanisms that drive motor rehabilitation through passive pneumotactile stimuli.

### 4.4 The role of right cerebellar lobule VI

The right cerebellar lobule VI has been suggested to be part of the sensorimotor somatotopic representations for the face (Grodd et al., 2001). In the present study, DCM results support that the right cerebellar lobule VI plays a role in the sensorimotor system during passive orofacial pneumotactile stimulation. Converging evidence from functional neuroimaging studies has shown that the cerebellum has multiple sensorimotor somatotopic representations (Bernard et al., 2013; Boillat, Bazin, & Van Der Zwaag, 2020; Bushara et al., 2001; Grodd et al., 2001; Kipping et al., 2013; Mottolese et al., 2013) and plays a critical role in motor functions (i.e., motor control, motor learning, etc.) and sensorimotor integration (Baumann et al., 2015; Buckner, Krienen, Castellanos, Diaz, & Yeo, 2011; Wolpert, Diedrichsen, & Flanagan, 2011). Our DCM results revealed effective connectivity of the cortico-cerebral networks and suggested the involvements of both forward (connections to the cerebellar) and backward (connections from the cerebellar) connections in cerebral-motor cortex connectivity during a passive pneumotactile stimuli stimulation paradigm. However, our DCM results were limited due to significant individual differences in cerebellar activation patterns. Only seven participants’ time series from 5 cm/s stimuli and five participants’ time series from 25 cm/s were successfully extracted and included in the DCM analyses. A recent study also reported that vibrotactile stimulation paradigms produced weaker activity in the cerebellum than the motor paradigms (Ashida, Cerminara, Edwards, Apps, & Brooks, 2019). Resting-state functional MRI data has reported the functional connectivity between the right lobule VI and the left M1 (Kipping et al., 2013). Our DCM results provided direct evidence on both forward and backward intrinsic connections between the right lobule VI and the contralateral left SI and M1 during the velocity pneumotactile stimuli. However, the modulation effect from the tactile stimulation was not significant, which may be due to the small number of participants included in the cerebellum DCM analysis or weak activity in the cerebellum during passive tactile stimulation paradigms.

## 5. CONCLUSIONS

In summary, the present study examined effective connectivity evoked by the orofacial pneumotactile perception of velocity using DCM on 20 neurotypical adults’ fMRI data, which has not been reported previously. Our DCM results demonstrated both similarities and differences in effective connectivity across the three-velocity orofacial pneumotactile stimulation. First, our DCM analyses suggested that the low velocity orofacial pneumotactile stimuli were processed in parallel through the contralateral SI and SII, and the medium and high-velocity stimuli recruited both serial and parallel processing types, supporting the coexistence of the parallel and serial processing theories during the passive orofacial pneumotactile stimulation paradigms. Second, the low and medium velocity orofacial pneumotactile stimuli modulated interhemispheric forward connections from the contralateral SII to the ipsilateral SII serially. Third, the significant modulation effect on forward connections between the contralateral SI and M1 during the low velocity orofacial pneumotactile stimulation supports the notion that the passive somatosensory inputs can affect the motor function. Therefore, the implication from our finding suggests that passive pneumotactile saltatory stimulation may bolster functional recovery during sensorimotor rehabilitation. Finally, we demonstrated that the right cerebellar lobule VI plays a role in the sensorimotor system during passive orofacial pneumotactile stimulation. In the future, we can design sensorimotor rehabilitation protocols for stroke survivors using multichannel TAC-Cell arrays and fMRI or MEG, or both to evaluate the efficacy of the rehabilitation protocols.

## Acknowledgements

We thank our participants and undergraduate research assistants from the UNL UCARE program, courtesy of the Pepsi Quasi Endowment and Union Bank & Trust.

